# Practice makes imperfect: stronger implicit interference with practice in individuals at high risk of developing Alzheimer’s disease

**DOI:** 10.1101/2023.05.16.541059

**Authors:** Shao-Min Hung, Sara W. Adams, Cathleen Molloy, Daw-An Wu, Shinsuke Shimojo, Xianghong Arakaki

## Abstract

Early screening to determine patient risk of developing Alzheimer’s will allow better interventions and planning but necessitates accessible methods such as behavioral biomarkers. Previously, we showed that cognitively healthy older individuals whose cerebrospinal fluid amyloid / tau ratio indicates high risk of cognitive decline experienced implicit interference during a high-effort task, signaling early changes in attention. To further investigate attention’s effect on implicit interference, we analyzed two experiments completed sequentially by the same high- and low-risk individuals. We hypothesized that if attention modulates interference, practice would affect the influence of implicit distractors. Indeed, while both groups experienced a strong practice effect, the association between practice and interference effects diverged between groups: stronger practice effects correlated with more implicit interference in high-risk participants, but less interference in low-risk individuals. Furthermore, low-risk individuals showed a positive correlation between implicit interference and EEG low-range alpha event-related desynchronization when switching from high-to low-load tasks. These results demonstrate how attention impacts implicit interference and highlight early differences in cognition between high- and low-risk individuals.

## 1. Introduction

Because current options for addressing Alzheimer’s disease are limited to preventative measures or palliative treatments, diagnostic methods that can screen for individuals at high risk of cognitive decline are critical for minimizing disease progression and allowing patients to make decisions about their future care while they are still cognitively healthy. Current methods of identifying such individuals are primarily limited to examining latent pathological changes such as levels of cerebrospinal fluid (CSF) biomarkers tau and amyloid beta (A*β*_42_). Although these biomarkers are well-established in the literature and may precede the onset of symptoms by over a decade [1], their potential use for widespread screening is limited by the expense and invasiveness of the lumbar puncture required for measurement. Behavioral biomarkers could bypass this problem, but introduce a new issue: given that most high-risk individuals are cognitively healthy prior to the onset of symptoms, it is extremely difficult to detect disease-dependent cognitive decline at the preclinical stage.

In a recent study, we investigated a possible solution by measuring how cognitively healthy high-risk and low-risk (defined by CSF A*β*_42_ / total tau protein ratio) older individuals responded to implicit distracting visual stimuli [2]. Motivated by studies that demonstrate immense interaction between attention and implicit processing in healthy young individuals [3,4], we hypothesized that early attentional changes could be revealed by evaluating how implicit information was processed in different individuals. Indeed, we found that cognitively healthy older individuals at high risk of cognitive decline experienced interference from an implicit (unseen) stimulus when the task difficulty was high. That is, participants’ reaction time slowed after an incongruent distractor was presented, even when they were not consciously aware of its existence. However, low-risk cognitively healthy older individuals were not susceptible to this implicit distraction. Furthermore, explicit distracting information (e.g., Stroop effect) interfered with the performance of both groups at a similar level. Taken together, these results suggest that early changes in implicit cognition are associated with the risk of cognitive decline, likely linked to Alzheimer’s.

Since attention deployment is at the core of implicit processing, here we further investigated whether *a reduction of task load due to practice* interacted with implicit interference. In a related study with healthy young participants, it was shown that short-term practice modulated the strength of implicit interference [4]. These findings provide a solid foundation for the current study, in which we examined two similar task-switching experiments [2,5] completed sequentially by the same cognitively healthy older individuals. The participants’ risk status for cognitive decline associated with Alzheimer’s disease had been previously determined for a longitudinal study and defined as cognitively healthy with normal A*β*_42_ /total tau protein ratio (CH-NATs, low-risk) or pathological ratio (CH-PATs, high-risk). We hypothesized that practice would protect the low-risk older participants from distraction: that is, a stronger practice effect would lead to weaker implicit interference by making it easier to suppress distracting information. In the high-risk older participants, we hypothesized that practice might pose no effect or even a reversed effect. Based on previous findings, high-risk older participants could be more likely to spend their attentional resources on distracting information. Thus, a reduction of task load could further strengthen implicit interference in this group.

## 2. Methods

### 2.1. Participants

Thirty-six cognitively healthy elderly (age range: 53-92) with no motor or vision difficulties were included from two prior studies [2,5]. These participants completed two separate experimental sessions with identical explicit tasks. Each session lasted approximately 50 minutes. The experimenters and the participants were oblivious to the physiological risk status of the participants. The institutional review boards (IRB) of the California Institute of Technology and the Huntington Medical Research Institutes (HMRI) approved this study (Quorum IRB, #27197). All participants gave written consent prior to participation. The final analysis included 17 CH-NATs and 19 CH-PATs (see Section 2.2) for behavioral analysis and 14 CH-NATs and 16 CH-PATs for EEG data analysis (6 were excluded because of artifacts). Participants’ demographics, including their age, sex, education year, and neuropsychological test scores, were reported in a previous study [2]. In each analysis, the demographics were comparable between the two groups (all *p* > .05).

### 2.2. Physiological status classification

A complete description of classification, including a list of neuropsychological testing, MRI diagnosis of small vessel disease, and the biochemical analysis of lumbar CSF, was detailed in several previous studies [5–7]. In short, only participants who were diagnosed without (1) cognitive impairment (Clinical Dementia Rating) or (2) psychiatric or neurological disorders were included in the current study. Different cognitive domains including memory, executive function, language, etc. were tested. The CSF samples were run on both Innotest (Innogenetics, discontinued) and MSD platforms (K15121G, MSD) to determine total tau and A*β*_42_. Participants were then classified depending on individual CSF A*β*_42_ / total tau ratios compared to a cutoff value (2.7132) derived from a logistic regression model that correctly diagnosed > 85% of clinically probable AD participants[6]. Amyloid / total tau ratios below the cutoff value classified participants as **c**ognitively **h**ealthy – **p**athological **a**myloid / **t**au, i.e., CH-PATs, while participants whose ratios were above the cutoff value were classified as **c**ognitively **h**ealthy – **n**ormal **a**myloid / **t**au, i.e., CH-NATs [2,5–7].

### 2.3. Experiment design

Participants completed both experiment sessions in the same quiet room. The visual stimuli were generated with E-Prime (Psychology Software Tools, Inc.) on a Dell Precision T5610 with a 20” screen. The participants’ movements were unconstrained, and they observed the visual stimuli from approximately 50 cm away.

In each trial, participants responded to two consecutive Stroop stimuli, i.e., words whose colors were incongruent with their literal meaning, such as the word RED written in green (Fig. 1). Participants were asked to name the color when the word was underlined (color-naming) and to name the word when the word was not underlined (word-naming). When the two stimuli prompted different tasks (e.g., word-naming for the first, and color-naming for the second), the trial was considered task-switching. Otherwise, if the tasks were the same, it was a non-task-switching trial. Participants were instructed to respond as accurately as possible and were given ample time for each response (up to 5 seconds). In the second experimental session, an additional gray word stimulus (17 ms) was added between the existing word stimuli with forward and backward masks (50 ms each). The masked word was either congruent or incongruent with the second word (target); congruent stimuli prompted the same response. However, no task was given to the masked word. Therefore, the explicit aspects of the tasks in each session were largely identical. More details of experimental design, rationale, and analysis are provided in the original study[2].

**Fig. 1.**
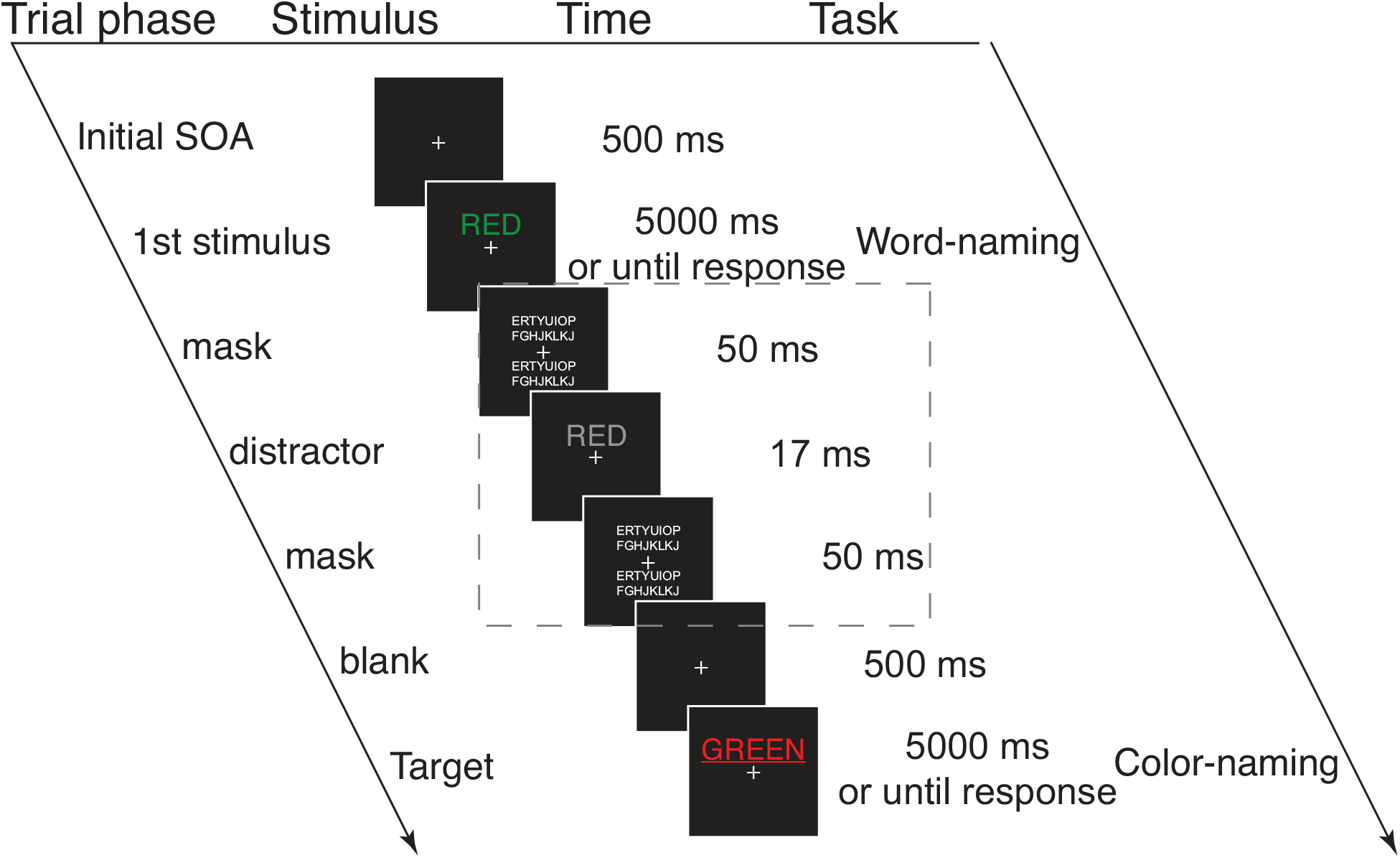
An example trial sequence (modified from [2] Fig. 1B). This is a task-switching trial (from word-naming to color-naming) with a congruent masked distractor. Each trial began with a 500-ms fixation, followed by the appearance of the first word stimulus for 5000 ms or until response. In the second experimental session only, a masked distractor (shown in the dashed box) was presented before the second word stimulus (i.e., target). The masked word was presented for 17 ms either above or below the fixation cross, sandwiched between two 50 ms masks consisting of random letters. No task was given to the masked distractor. After the first stimulus (experiment one) or distractor (experiment two), the screen was blank for 500 ms before the target was displayed for another 5000 ms or until response. The only difference between the two experiments was the masked distractor presentation. SOA: stimulus onset asynchrony.

At the start of each session, participants underwent a short period of practice until they felt comfortable proceeding to the main experiment. The experiments consisted of three blocks of 64 trials. In the second session, a surprise post-experiment awareness test was performed to assess the invisibility of the masked words. During this test, participants were asked to judge the location (top/bottom) of the masked stimulus in a two-alternative forced-choice manner. Chance performance on this test indicated a lack of conscious awareness of the distractor.

## 3. Results

### 3.1. Original key findings replicated: stronger implicit interference in CH-PATs

The analyses below are reported in the format of the mean (standard error of the mean [SEM]) unless otherwise specified. The implicit interference was deemed the target reaction time slowing caused by an incongruent implicit distractor, as compared to a congruent implicit prime. Because only 36 of 40 participants from the previous study were considered here, we repeated the same analyses to ensure that the results were similar. Indeed, the mean accuracy of the awareness test was 44.58% (2.32%) and slightly below chance (compared to 50%, *t*(29) = -2.33, *p* = 0.03), confirming the implicit nature of the distractor. No difference in distractor awareness was found between CH-NATs (44.26 (3.8) %) and CH-PATs (45.00 (2.21) %) (two-sample t-test, *t*(28) = 0.1543, *p* = 0.88). Similarly, we reproduced the key finding that CH-PATs had more interference from an implicit response-incongruent distractor than CH-NATs (CH-PATs: 4.21 (1.51) % change in reaction time, *t*(18) = 2.79, *p* = 0.01; CH-NATs: -0.31 (1.1) % change in reaction time, *t*(16) = -0.28, *p* = 0.78. A direct comparison between the two yielded *t*(34) = 2.37, *p* = 0.02, Cohen’s *d* = 0.81).

### 3.2. Practice has distinct effects on CH-PATs and CH-NATs

Both groups exhibited a practice effect in the second session. Overall, between the two sessions, the average reaction time decreased by 6.19% (3.04%), while the average accuracy increased by 6.09% (1.40%). The time interval between the two sessions was comparable across the two groups (CH-PATs: 15.84 (2.87) days; CH-NATs: 15.29 (1.68) days, *t*(34) = 0.16, *p* = 0.87).

We investigated whether practice over time differentially affects performance of the low-risk and high-risk participants. To this end, we first performed two separate mixed-effect analyses of variance (ANOVAs) on the mean accuracy and on the reaction time with one between-subject factor (participant status: CH-NATs / CH-PATs) and one within-subject factor (session number: one / two). Analysis of the accuracy yielded a main effect of session number, *F*(1, 34) = 21.66, *p* < 0.0001, *ηp2* = 0.39, and an interaction between session and participants’ status, *F*(1, 34) = 9.03, *p* = 0.005, *ηp2* = 0.21. The analysis of reaction time yielded a main effect of session, *F*(1, 34) = 4.20, *p* = 0.048, *ηp2* = 0.11. Post-hoc analyses revealed that CH-PATs exhibited a stronger practice effect in the accuracy (CH-NATs: 2.08% (1.36%) vs. CH-PATs: 9.67% (2.05 %), *t*(34) = -3.01, *p* = 0.005) but not reaction time (CH-NATs: 4.06% (4.44%) vs. CH-PATs: 8.10% (4.24%), *t*(34) = -0.66, *p* = 0.52).

To address the impact of practice on implicit interference, we ran a correlation analysis between the implicit distractor interference effect and the practice effect in CH-NATs and CH-PATs, respectively. For the purpose of this analysis, the practice effect was defined as the percentage decrease of reaction time in the second session compared to the first because the interference effect was only observed in the domain of reaction time. By converting reaction times into percentages for both interference and practice effects, baseline performance differences and the long tails in typical reaction time distributions were eliminated. Analysis of the CH-NATs yielded a marginal negative correlation between practice and implicit interference (*r* = -0.46, *p* = 0.06). However, the same analysis on CH-PATs yielded a positive correlation between the two (*r* = 0.50, *p* = 0.03). We calculated z-scores for the r values and directly compared the two groups’ correlations with a two-tailed test, which resulted in a p-value of 0.004. These results showed that the interaction between practice and the implicit interference was distinct between CH-NATs and CH-PATs: better performance in the second session was correlated with less implicit distraction in CH-NATs, while for CH-PATs, better performance in the second session led to stronger distraction (Fig. 2., left).

**Fig. 2.**
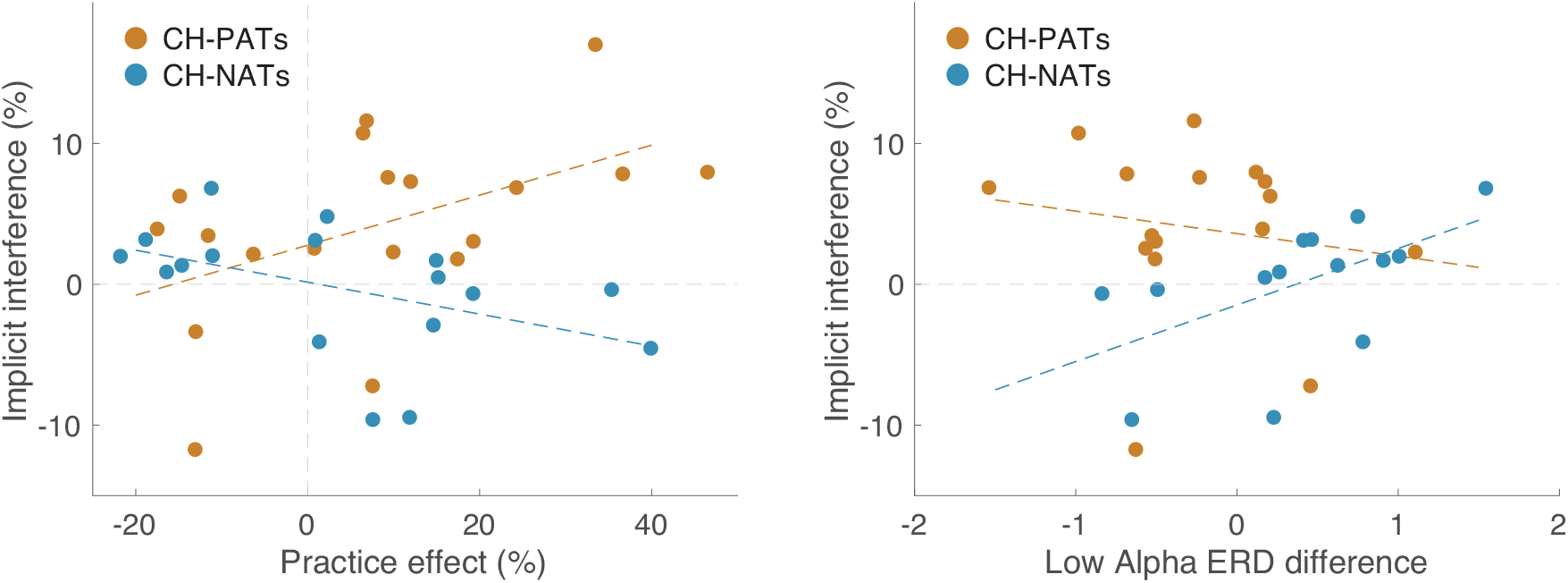
Left: Distinct correlations between practice effect (percentage decrease of reaction time in the second session; positive values indicate a stronger practice effect, i.e. more decrease in reaction time, x-axis) and implicit interference (percentage decrease of reaction time between incongruent distractor and congruent prime; positive values indicate stronger interference, i.e. more increase in reaction time, y-axis) in CH-NATs (low-risk) and CH-PATs (high-risk). Right: Positive correlation between lower alpha (8-11 Hz) ERD (event-related desynchronization) difference (x-axis) and the implicit interference (y-axis) was observed only in CH-NATs. Each dot represents a participant. The dotted lines were linearly fitted to the two correlations.

### 3.3. Greater low-range alpha ERD difference between task-switching and non-switching trials on EEG in session one is correlated with interference in CH-NATs

In the first session, EEG data was collected for the majority of participants (14 of 17 CH-NATS and 16 of 19 CH-PATs), which allowed us to determine the correlation between prior EEG data and later behavioral performance in these two groups. To minimize multiple comparisons and to focus on an attention component, we examined the most distinctive alpha event-related desynchronization (ERD) difference between the CH-NATs and CH-PATs. In the original study, we calculated the difference in the central area (C3, Cz, C4) alpha ERD between high-load task-switching trials and low-load non-switching trials. For CH-PATs, this difference was negative, indicating that these participants had stronger low alpha ERDs during switching trials. For CH-NATs, the opposite was observed: their alpha ERD difference was positive, indicating a stronger low alpha ERD during non-switching trials. Together, these results suggested that CH-PATs have stronger attentional engagement during task switching, while CH-NATs have higher attentional engagement when not changing tasks. These results are also found with the reduced cohort in our study (low alpha ERD mean difference (SD), CH-PATs: -0.26 (0.63) vs. CH-NATs: 0.37 (0.67), *t*(28) = -2.68, *p* = 0.01, Cohen’s *d* = 1.00).

We ran a correlation analysis between the aforementioned low alpha ERD difference in the first session and the implicit interference in the second session in both groups. CH-PATs had no correlation between their low alpha ERD difference and implicit interference (*r* = -0.16, *p* = 0.54). In contrast, there was a positive correlation between low alpha ERD difference and implicit interference in CH-NATs (*r* = 0.56, *p* = 0.04) (Fig. 2., right). A further z-score comparison between the two correlations revealed a p-value of 0.05. These results showed that CH-NATs who exhibited a higher low-range alpha ERD difference in the first session also experienced stronger distraction in the second session. This finding suggests that in low-risk participants, having lower attentional engagement in switching trials is associated with stronger implicit interference.

## 4. Discussion

In our two original studies, we examined the electrophysiological signatures [5] during a cognitively demanding task as well as the behavioral response to implicit distracting information [2] in cognitively healthy older individuals with a high risk (CH-PATs) or low risk (CH-NATs) of cognitive decline. The largely identical experimental designs and participants of these studies created a unique opportunity for the current study to investigate if practice or the EEG signatures found in the first session were associated with implicit interference. Overall, we found that practice had distinct effects on high-risk and low-risk individuals. While the practice effect was negatively correlated with interference in the low-risk participants, a positive correlation was found in the high-risk participants. Furthermore, in low-risk participants, stronger interference was associated with a greater low-range alpha ERD difference between task-switching and non-switching trials, suggesting that lower attentional engagement in switching trials is associated with susceptibility to interference. Although the low-risk participants had similar performance to what has been observed in a healthy young population [4], our study provides evidence that an increase in implicit interference with decreased task load is not a part of normal aging. Instead, these changes provide a potential indicator for risk of cognitive decline.

The opposite correlations between practice and implicit interference in the low-risk and high-risk populations indicate distinct underlying cognitive infrastructures in these two populations. If practice reduced cognitive load, the participants should have more cognitive resources available in the second session. Based on our results, low-risk and high-risk participants utilized these additional cognitive resources differently. Low-risk individuals learned to better inhibit distracting implicit information, while high-risk individuals deployed additional attentional resources to the distracting stimulus. This could explain why a stronger alpha ERD was observed in high-risk older individuals during various challenging tasks [5,7,8], including the first session of our study. That is, high-risk individuals tended to deploy additional attentional resources to both the target and the distracting feature. This approach could potentially maintain participant performance on the target at the expense of processing additional irrelevant sensory information, which kept the high-risk individuals overtly cognitively healthy. Future research is needed to unravel the neural machinery of the attention network in high-risk individuals.

In low-risk individuals, the positive correlation between low alpha ERD difference between task-switching and non-switching trials in the first session and implicit interference in the second session illustrates the similarities between low-risk cognitively healthy older adults and healthy young participants. That is, when low-risk participants are more attentive to a target, implicit interference is less likely to occur, just as in younger individuals. Recently, with concurrent tracking of target- and distractor-triggered alpha oscillations, Gutteling and colleagues [9] showed that distractor-triggered alpha oscillations increased significantly when the target was more difficult to process. These results from young individuals offer another explanation regarding distractors: task load could modulate alpha power in response to both distractors and targets. Future research on older individuals is needed to disentangle the target and distractor effects and to strengthen the current results with a larger sample size. Taken together, however, our results provide evidence that practice, although typically considered a positive effect, may uniquely influence high-risk individuals by increasing susceptibility to implicit interference. In low-risk participants, however, practice appears to remain positive as their behavioral and neurophysiological data demonstrate high similarity with that of younger individuals. How and when the practice effects of the two groups bifurcate is a critical research question for better understanding early cognitive decline in Alzheimer’s.

## Acknowledgments

The current study was not preregistered. We are grateful for the research funding provided by the James Boswell Postdoctoral Fellowship, the Caltech BBE Divisional Postdoctoral Fellowship, and the sub-award under the Aligning Consciousness Research with US Funding Mechanisms by Templeton World Charity Foundation (TWCF: 0495) to S.-M.H., L. K. Whittier Foundation to HMRI, and the National Institute of Health R56 (R56AG063857) and R01 (R01AG063857) to S.-M.H, D.-A. W., S.S., and X.A.

## Data availability

The raw data and codes are available upon reasonable request.

## References

1 Mattsson-Carlgren N, Leuzy A, Janelidze S, Palmqvist S, Stomrud E, Strandberg O, et al. The implications of different approaches to define AT(N) in Alzheimer disease. Neurology. 2020 May;94(21):e2233–44.

2 Hung S, Wu D, Shimojo S, Arakaki X. Stronger implicit interference in cognitively healthy older participants with higher risk of Alzheimer’s disease. Alz & Dem Diag Ass & Dis Mo. 2022 Jan;14(1). DOI: 10.1002/dad2.12340

3 Kiefer M, Martens U. Attentional sensitization of unconscious cognition: Task sets modulate subsequent masked semantic priming. Journal of Experimental Psychology: General. 2010;139(3):464–89.

4 Hung S-M, Wu D-A, Shimojo S. Task-induced attention load guides and gates unconscious semantic interference. Nature Communications. 2020 Apr;11(1):2088.

5 Arechavala RJ, Rochart R, Kloner RA, Liu A, Wu D-A, Hung S-M, et al. Task switching reveals abnormal brain-heart electrophysiological signatures in cognitively healthy individuals with abnormal CSF amyloid/tau, a pilot study. International Journal of Psychophysiology. 2021 Dec;170:102–11.

6 Harrington MG, Chiang J, Pogoda JM, Gomez M, Thomas K, Marion SD, et al. Executive Function Changes before Memory in Preclinical Alzheimer’s Pathology: A Prospective, Cross-Sectional, Case Control Study. PLoS ONE. 2013 Nov;8(11):e79378.

7 Arakaki X, Hung S-M, Rochart R, Fonteh AN, Harrington MG. Alpha desynchronization during Stroop test unmasks cognitively healthy individuals with abnormal CSF Amyloid/Tau. Neurobiology of Aging. 2022 Apr;112:87–101.

8 Arakaki X, Lee R, King KS, Fonteh AN, Harrington MG. Alpha desynchronization during simple working memory unmasks pathological aging in cognitively healthy individuals. PLoS ONE. 2019 Jan;14(1):e0208517.

9 Gutteling TP, Sillekens L, Lavie N, Jensen O. Alpha oscillations reflect suppression of distractors with increased perceptual load. Progress in Neurobiology. 2022 Jul;214:102285.

